# The yeast Gcn2 protein kinase mediates P-body formation both through its activity with eIF2α and with non-canonical substrates

**DOI:** 10.64898/2026.11.05.764498

**Authors:** Viktoras Stonys, Simon J. Hubbard, Mark P. Ashe, Chris M. Grant

## Abstract

P-bodies (PBs) are control hubs for RNA processing and storage that form when translation initiation is inhibited, including during stress. In yeast, Gcn2 is the sole eIF2α kinase that inhibits translation initiation in response to diverse stresses. We show that PB formation depends strongly on Gcn2 during amino acid starvation but only partially during oxidative stress. Constitutively active Gcn2 also induces PB formation in unstressed cells. Surprisingly, PBs are still constitutively formed in a *gcn3* mutant strain that is insensitive to eIF2α phosphorylation suggesting that Gcn2 also promotes PB formation independently of eIF2α phosphorylation. Analysis of several putative Gcn2-substrates involved in RNA metabolism identified several proteins required for Gcn2-mediated PB formation. Among these, Smu2, which was not previously implicated in PB formation, localises to PBs and is required for stress-induced coalescence of mRNA granules into PBs. Together, our findings indicate that Gcn2 promotes PB formation via two complementary routes: eIF2α phosphorylation, which generates untranslated mRNAs, and Smu2-dependent maturation of mRNA granules into PBs.

**Summary statement:** The Gcn2 protein kinase pays a key role in P-body formation during stress conditions that is not solely dependent on its canonical eIF2α substrate

## Introduction

All cells are required to maintain proteostasis in the face of fluctuating cellular demands, particularly under conditions of stress. To achieve this, cells have evolved regulatory pathways that attenuate global protein synthesis while selectively reprogramming translation to support stress-specific, pro-survival responses (Clemens, 2001, Proud, 2005). A highly conserved mechanism underlying this form of translational control is phosphorylation of the α subunit of eukaryotic initiation factor 2 (eIF2α) by the protein kinase Gcn2, which suppresses overall protein synthesis while preferentially enhancing the translation of stress-responsive genes that promote cellular adaptation and survival (Wek, 2018).

Phosphorylated eIF2α inhibits the guanine nucleotide exchange factor (GEF) activity of eIF2B, thereby impairing the regeneration of eIF2–GTP required for translation initiation (Dever et al., 2016, Krishnamoorthy et al., 2001, Sudhakar et al., 2000). The resulting depletion of the eIF2–GTP pool restricts formation of the ternary complex composed of eIF2–GTP and initiator methionyl tRNA, resulting in compromised translation initiation and delayed ribosomal reinitiation (Hinnebusch, 2005). The reduced reinitiation efficiency selectively enhances the translation of mRNAs critical for stress adaptation, including *GCN4* in yeast and *ATF4* in mammals (Dever et al., 1992, Lu et al., 2004, Vattem and Wek, 2004). Gcn4 and ATF4 are transcription factors that reprogram gene expression to promote cellular adaptation by regulating amino acid metabolism, redox homeostasis, mitochondrial function, autophagy and other pro-survival pathways under conditions of stress (Iurlaro et al., 2017, Natarajan et al., 2001, Shenton et al., 2006, Siu et al., 2002, Wang et al., 2010). In mammalian cells, four distinct eIF2α kinases have been identified which, despite responding to diverse upstream stress signals, converge on eIF2α phosphorylation and are therefore collectively defined as core components of the integrated stress response (ISR) pathway (Donnelly et al., 2013, Wek, 2018).

In yeast, Gcn2 is the sole eIF2α kinase, which phosphorylates eIF2α in response to various nutrient depletion conditions and exposure to diverse stress conditions (Hinnebusch and Fink, 1983, Castilho et al., 2014, Masson, 2019, Gupta and Hinnebusch, 2023). These different growth conditions are thought to activate Gcn2 via an accumulation of deacylated tRNAs that bind to the histidyl-tRNA synthetase (HisRS)-like domain of Gcn2 on ribosomes (Wek et al., 1995, Lageix et al., 2015). However, many conditions that are not associated with the accumulation of deacylated tRNAs activate Gcn2 and recent studies have identified collided ribosomes as critical activators of Gcn2 (Masson, 2019, Gupta and Hinnebusch, 2023, Nanjaraj Urs et al., 2024, Nanjaraj Urs et al., 2025, Chang and Choe, 2026). The Gcn1 and Gcn20 proteins have been identified as essential upstream regulators that facilitate activation of Gcn2 during amino acid starvation conditions (Marton et al., 1997, Sattlegger and Hinnebusch, 2005). Gcn1 is a key adapter protein that binds both Gcn2 and the ribosome. It is required for eIF2α phosphorylation in response to amino acid starvation (Marton et al., 1993, Sattlegger and Hinnebusch, 2000). Additionally, full Gcn2 activation requires Gcn20 that stabilizes the interaction of Gcn1 with ribosomes (Marton et al., 1997, Pochopien et al., 2021). This has led to a model where Gcn1 and Gcn20 enable Gcn2 activation by coupling uncharged tRNAs to ribosome-bound Gcn2 (Marton et al., 1997). Consistent with the idea that collided ribosomes activate the ISR, Gcn1 binds disomes that are formed by collisions between stalled ribosomes and trailing elongating ribosomes (Pochopien et al., 2021, Oltion et al., 2023). These findings suggest that Gcn1, Gcn20, and Gcn2 act at collided ribosomes to integrate signals from translational stress and uncharged tRNAs, thereby ensuring context-specific activation of the ISR.

Alongside translational inhibition and reprograming during stress, mRNAs can be sequestered into ribonucleoprotein (RNP) granules that play key roles in RNA metabolism. Prominent examples include processing bodies (PBs), cytoplasmic granules that act as sites of mRNA storage and decay (Jain and Parker, 2013, Luo et al., 2018, Standart and Weil, 2018, Feng et al., 2026). PB formation is triggered by conditions that increase the pool of translationally repressed mRNAs, causing non-translating mRNAs to accumulate and become sequestered away from the actively translated pool (Teixeira et al., 2005, Teixeira and Parker, 2007, Lui et al., 2014). This interdependence of PB formation and repression of translation ensures that mRNA metabolism and protein synthesis are coordinated processes, enabling cells to rapidly adapt to changing conditions (Kedersha et al., 2005, Standart and Weil, 2018). Surprisingly therefore, little is known regarding the role of Gcn2-mediated translation initiation inhibition in PB formation. Studies in mammalian cells have shown PBs are still formed in cells in the absence of eIF2α phosphorylation suggesting that eIF2α phosphorylation is not essential for PB formation (Kedersha et al., 2005). In yeast cells, the presence of a constitutively active Gcn2 allele was found to enhance the formation of PBs during glucose starvation (Buchan et al., 2008), whereas another study found no correlation between eIF2α phosphorylation and PB formation (Hoyle et al., 2007). Thus, the full extent of how or whether Gcn2-mediated eIF2α phosphorylation contributes to PB formation is unclear.

Here, we examined the roles of Gcn2 and eIF2α phosphorylation in PB formation under three stresses: amino acid starvation, glucose starvation and oxidative stress, that all promote PBs. Gcn2 was dispensable for PB formation during glucose starvation, which does not promote eIF2α phosphorylation. In contrast, Gcn2 plays a major role in PB formation during amino acid starvation conditions and a more limited role in PB formation during oxidative stress conditions, despite both conditions promoting eIF2α phosphorylation. Surprisingly however, PB formation depended only partially on the canonical function of Gcn2 in phosphorylating eIF2α and we have identified several putative Gcn2-substrate proteins that mediate Gcn2-dependent PB formation during stress conditions.

## Results

### Diverse stress conditions that inhibit translation initiation promote similar PB formation

PB formation depends on the dynamic interplay between translational activity and mRNA metabolism. It is intrinsically linked to the repression of translation initiation where ribosome-free mRNAs accumulate and become sequestered in PBs away from the actively translated pool of cellular mRNAs (Teixeira et al., 2005, Teixeira and Parker, 2007). We investigated PB formation following three different stresses known to cause an inhibition of translation initiation: amino acid starvation, glucose deprivation and H_2_O_2_-induced oxidative stress (Ashe et al., 2000, Shenton et al., 2006, Hoyle et al., 2007). Glucose starvation and oxidative stress have been shown to promote PB formation in yeast (Teixeira et al., 2005, Buchan et al., 2008), whereas amino starvation is not thought to promote PB formation despite causing an inhibition of translation initiation (Hoyle et al., 2007).

We selected Dcp1 as a well-established PB marker protein based on its extensive use in previous studies (Sheth and Parker, 2003, Hoyle et al., 2007, Buchan et al., 2008). Using live-cell fluorescence microscopy, we examined Dcp1-GFP localisation in unstressed cells and after one hour of glucose starvation, amino acid starvation and exposure to hydrogen peroxide (Fig. 1A). Diffuse Dcp1-GFP fluorescence was observed in most cells during non-stress conditions, whereas all three stresses induced the formation of multiple Dcp1-GFP granules. A previous study found no PB formation after shifting cells from complete SCD media to media lacking amino acids for 30 mins (Hoyle et al., 2007). The longer, one hour starvation period used in our experiments may therefore account for the Dcp1 granule formation observed in our experiments.

**Figure 1.**
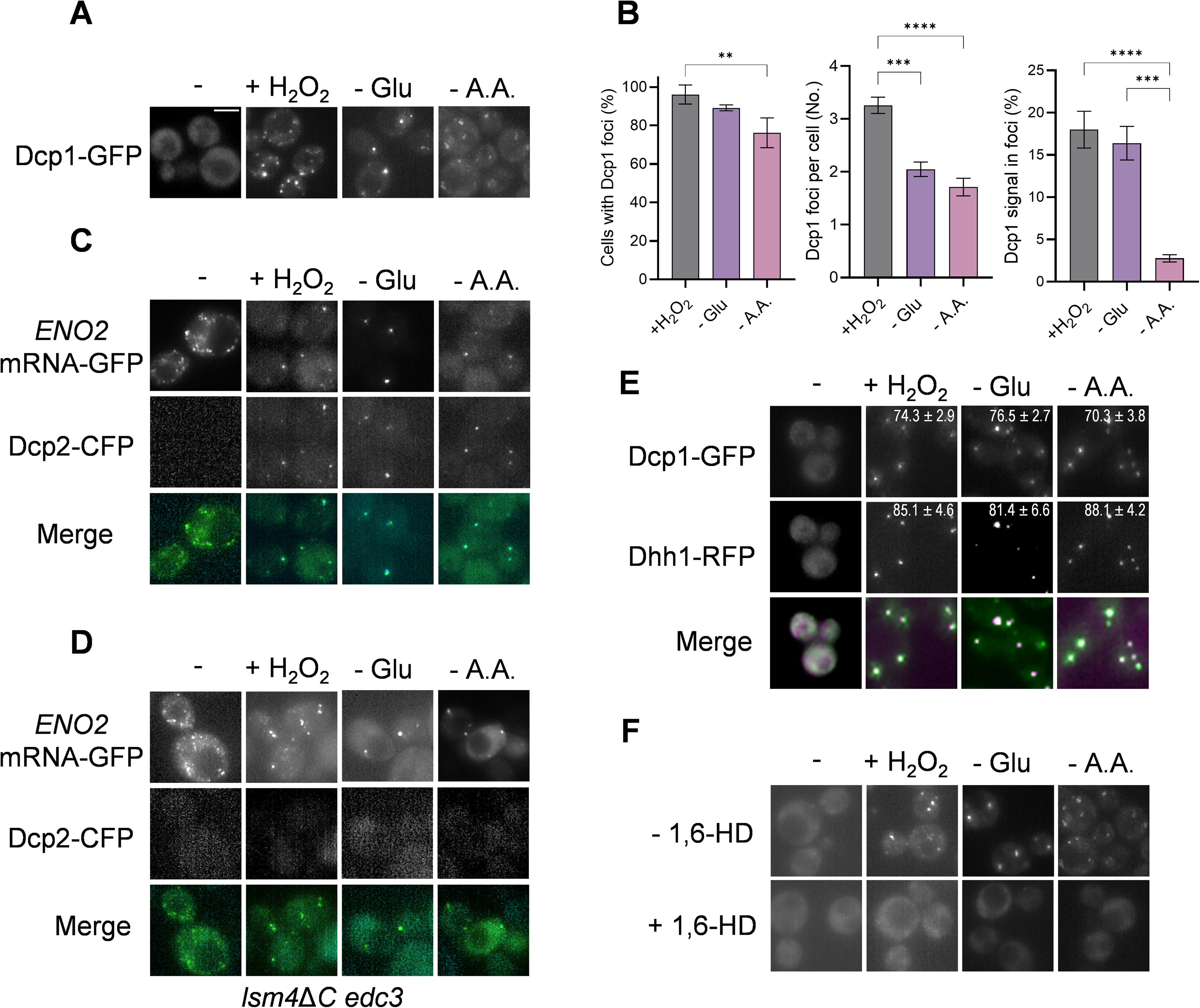
Stress conditions that inhibit translation initiation promote similar PB formation. **A.** Representative epifluorescent microscopic images are shown for strains expressing Dcp1-GFP. Strains were grown to exponential phase and left untreated (-), starved for glucose (-Glu), starved for ammino acids (-AA) or treated with 0.8 mM hydrogen peroxide (+H_2_O_2_) all for one hour. Scale bar 4 µm. **B.** Charts show the percentage of cells containing Dcp1-GFP foci, the average number of Dcp1-GFP foci per cell and the percentage Dcp1-GFP signal localised in foci (*n* > 100, based on three biological replicates). Error bars denote standard deviation. Significance is shown using a one-way ANOVA test; ** < 0.01, *** < 0.005, **** < 0.001. **C.** *ENO2*-MS2 mRNA was visualised in wild-type cells expressing GFP-fused coat protein and endogenously tagged Dcp2-CFP. **D.** As for (C) but in an *lsm4*Δ*C edc3*Δ mutant. **E.** PB formation is shown for strains expressing Dcp1-GFP and Dhh1-RFP. Dhh1-RFP is shown in magenta. Numbers indicate the percentage of all Dcp1 or Dhh1 foci co-localised with either of the PB marker proteins (*n* > 100, based on three biological replicates). **F.** Live cell imaging of cells described in (A) treated with 5% 1,6-hexanediol for 2 min to dissolve phase-separated condensates.

We found that at least 90% of cells contained Dcp1 granules when subjected to hydrogen peroxide stress or glucose starvation, whereas amino acid starvation-induced Dcp1 granules were visible in ∼75% of cells (Fig. 1B). On average, around three Dcp1 granules per cell were formed under oxidative stress, compared with up to two granules following glucose or amino acid starvation (Fig. 1B). The Dcp1 granules formed following amino acid starvation were noticeably fainter than those formed following oxidative stress or glucose starvation conditions (Fig. 1A). Quantification of the cellular Dcp1-GFP signal revealed that 15-17% of Dcp1-GFP was localized to granules under oxidative stress or glucose starvation conditions, compared with 2.5% of Dcp1-GFP following amino acid starvation (Fig. 1B).

The localisation of Dcp1-GFP to granules is not sufficient to define PB formation since Dcp1 may localize to other intracellular sites such as non-specific protein aggregates. We therefore examined four further characteristics of PBs, primarily defined under glucose starvation. Firstly, we examined exemplar mRNA localization following our different stress conditions, given that RNP granules such as PBs contain certain mRNAs. For example, the *ENO2* mRNA, encoding enolase, has previously been shown to localise to translation factories termed CoFe granules in unstressed cells, then following glucose depletion the *ENO2* mRNA granules rapidly coalesce and recruit mRNA decay components (Lui et al., 2014, Morales-Polanco et al., 2021). For these experiments, we visualized *ENO2* using the m-TAG system, where the MS2 bacteriophage system is used to tether GFP to an mRNA to study its localization in live cells (Haim-Vilmovsky et al., 2011). *ENO2* mRNA was present in multiple granules in unstressed cells which coalesced into fewer, brighter granules in cells subjected to our three stress conditions, consistent with *ENO2* mRNA entering PBs (Fig. 1C). In agreement with previous observations (Lui et al., 2014), *ENO2* mRNA localized with the Dcp2-CFP granules formed in response to glucose depletion. Similar co-localization was also observed between *ENO2* mRNA and Dcp2-CFP following amino acid starvation or oxidative stress conditions (Fig. 1C).

Secondly, we examined an *edc3*Δ *lsm4*Δ*C* mutant which lacks the Edc3 decapping activator and contains a mutant version of Lsm4 where its prion-like domain has been deleted. This mutant is defective in PB formation induced by glucose depletion (Decker et al., 2007). Whilst PB formation is suppressed in *edc3*Δ *lsm4*Δ*C* mutants, mRNAs still condense into granules following glucose depletion, albeit at a reduced frequency (Lui et al., 2014). We found that Dcp2-CFP granule formation was similarly suppressed following oxidative stress and amino acid starvation consistent with this mutant suppressing PB formation in all three of our stress conditions (Fig. 1D).

Thirdly, we characterized the localization of Dhh1, another PB marker protein, during our different stress conditions. Dhh1 is an mRNA decapping activator that has been extensively localized to glucose-starvation-induced PBs (Swisher and Parker, 2010, Cary et al., 2015, Kershaw et al., 2021). Similar to Dcp1, diffuse Dhh1-RFP fluorescence was observed in most cells during non-stress conditions, and multiple Dhh1-RFP granules were formed in response to all three stress conditions (Fig. 1E). Greater than 70% of the Dhh1 granules were co-localised with Dcp1 following all three stress conditions.

Finally, we treated cells with 1,6-Hexanediol which disrupts the weak hydrophobic interactions that drive liquid-liquid phase separation and rapidly dissolves condensates such as PBs in yeast and mammalian cells (Kroschwald et al., 2015, Wheeler et al., 2016). Treatment with 1,6-Hexanediol for two minutes resulted in the complete loss of the Dcp1 granules formed under all three stress conditions (Fig. 1F). Taken together, our data indicate that similar PBs are induced following glucose starvation, oxidative stress and amino acid starvation.

### Gcn2 activity is required for stress-induced PB formation

Given the established link between translational repression and PB formation, we examined whether the Gcn2 kinase is required for PB formation under stresses that inhibit translation initiation through different mechanisms. Amino acid starvation and oxidative stress activate Gcn2, leading to eIF2α phosphorylation and inhibition of translation initiation (Hinnebusch, 2005, Shenton et al., 2006). In contrast, glucose starvation rapidly inhibits translation initiation independently of eIF2α-P formation by disrupting the eIF4A interaction with the 48S preinitiation complex (Ashe et al., 2000, Castelli et al., 2011, Bresson et al., 2020). Prolonged glucose starvation (six hours) has also been shown to increase eIF2α-P levels (Yang et al., 2000), whereas eIF2α phosphorylation is unaffected by the shorter periods of glucose starvation typically used to induce PB formation (Hoyle et al., 2007).

We first confirmed that hydrogen peroxide treatment and amino acid starvation induce eIF2α-P formation under our experimental conditions (Fig. 2A). This phosphorylation depended on Gcn2 since it was undetected in a *gcn2* mutant and was restored via complementation from plasmid-based Gcn2 expression (Fig. 2A). As expected, no phosphorylation of eIF2 was detected following glucose starvation imposed for one hour (Supplementary Fig. 1A). Translational activity was analysed by examining the sedimentation of ribosomes within a sucrose density gradient during non-stress and stressed conditions. Amino acid starvation, glucose deprivation and H_2_O_2_-induced oxidative all caused a marked inhibition of translation initiation as expected (Ashe et al., 2000, Shenton et al., 2006), shown by an increase in the 80S monosome peak and reduced levels of actively translating polysomes (Supplementary Fig. 1B). This inhibition was largely dependent on Gcn2 during amino acid starvation and oxidative stress but Gcn2-independent during glucose starvation-induced inhibition (Supplementary Fig. 1B).

**Figure 2.**
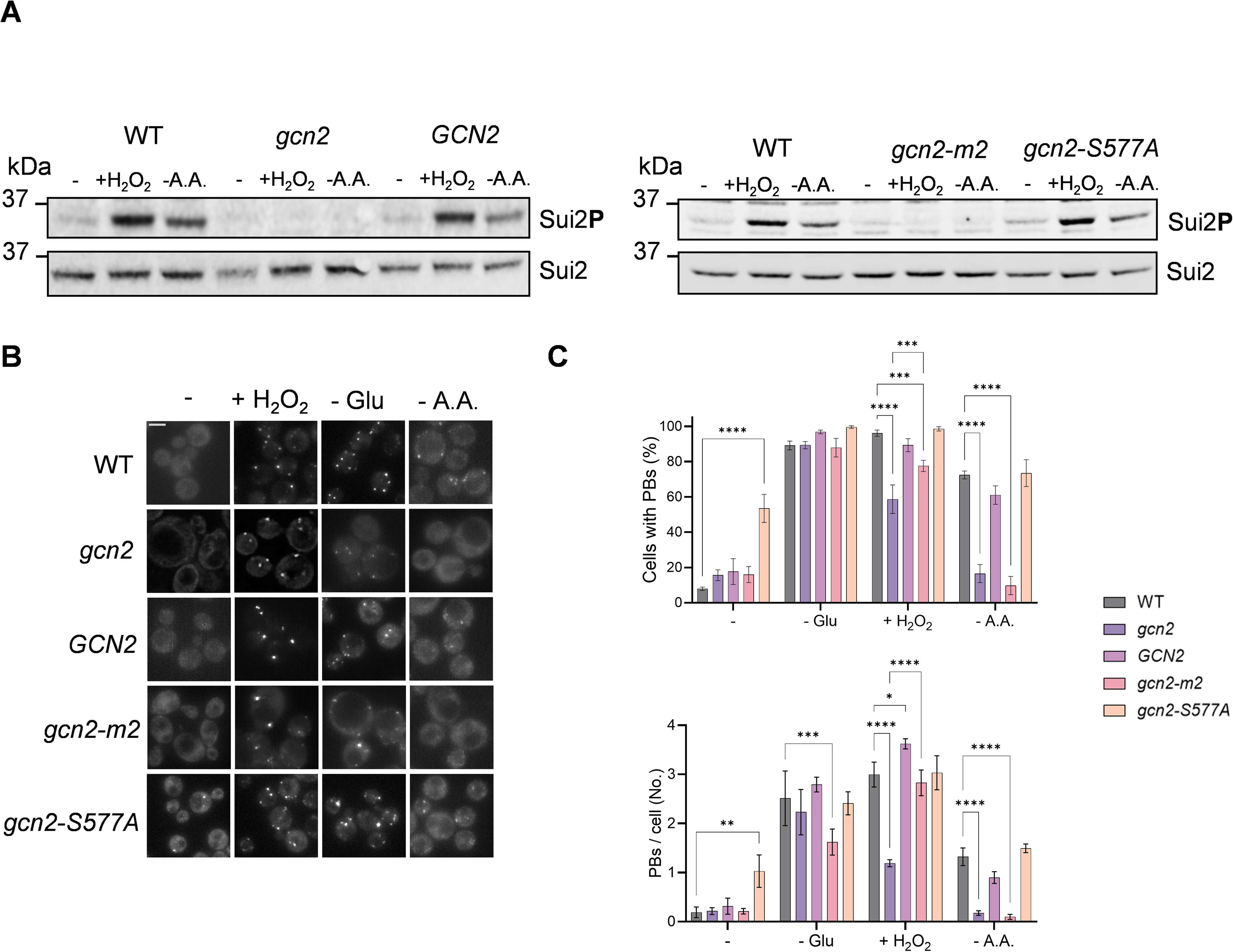
Requirement for Gcn2 activity during stress-induced PB formation. **A.** Phosphorylation of eIF2α is shown for strains grown to exponential phase and left untreated (-), starved for glucose (-Glu), starved for ammino acids (-AA) or treated with hydrogen peroxide (+H_2_O_2_). Whole cell extracts were prepared from wild-type strains containing vector (WT) and *gcn2* mutant strains containing vector (*gcn2*) or vectors expressing *GCN2, gcn2-m2 gcn2-S577A.* Western blots are shown probed with antibodies that recognise eIF2α-P (Sui2P) or eIF2α (Sui2) as a loading control. **B.** Representative epifluorescent microscopic images are shown for strains expressing Dcp1-GFP grown as above. Scale bar 4 µm. **C.** Charts show the percentage of cells containing Dcp1-GFP foci and the average number of Dcp1-GFP foci per cell (*n* > 100, based on three biological replicates). Error bars denote standard deviation. Significance is shown using a one-way ANOVA test; ** < 0.01, *** < 0.005, **** < 0.001.

PB formation was again monitored via localisation of Dcp1-GFP using live-cell fluorescence microscopy in wild-type and *gcn2* mutant strains (Fig. 2B). Under oxidative stress, Gcn2 was partially required for PB formation, since the number of cells with PBs and the average number of PBs per cell was significantly reduced in the *gcn2* mutant strain (Fig. 2C). In contrast, PB formation was almost completely abrogated in the *gcn2* mutant following amino acid starvation conditions (Fig. 2C). Restoring *GCN2* expression from a plasmid complemented the defect in PB formation for both oxidative stress and amino acid starvation. No requirement for Gcn2 was observed for PB formation following glucose depletion (Fig. 2C).

Gcn2 activation involves uncharged tRNA binding to its histidyl-tRNA synthetase-related domain, which triggers kinase autophosphorylation (Qiu et al., 2002). Gcn2 activity is also regulated by Tor-dependent phosphorylation of Ser577 (Cherkasova and Hinnebusch, 2003). To investigate how these regulatory mechanisms affect PB formation, we examined two Gcn2 mutants: *gcn2-m2*, which abolishes uncharged tRNA binding (Dong et al., 2000), and *gcn2-S577A*, which increases affinity for uncharged tRNAs and causes constitutive eIF2α phosphorylation in unstressed cells (Cherkasova and Hinnebusch, 2003). We first confirmed the expected effects on eIF2α phosphorylation: phosphorylation was abolished in the *gcn2-m2* mutant but was somewhat constitutive in the *gcn2-S577A* mutant (Fig. 2B).

Consistent with its constitutive eIF2α phosphorylation, the *gcn2-S577A* mutant formed PBs even in the absence of stress (Fig. 2C). By contrast, amino acid starvation-induced PB formation was almost completely abolished in the *gcn2-m2* mutant, as in the *gcn2* deletion mutant. This finding supports a model in which uncharged tRNA binding activates Gcn2 to promote PB formation during amino acid starvation. Under oxidative stress, however, PB formation was greater in the *gcn2-m2* mutant than in the *gcn2* deletion mutant, despite the absence of eIF2α phosphorylation in *gcn2-m2* cells. Together, these results indicate that Gcn2 is essential for PB formation during amino acid starvation but contributes only partially during oxidative stress.

### An eIF2α-P-independent role for Gcn2 in PB formation

We next examined whether Gcn2-dependent PB formation is entirely dependent on the phosphorylation of its canonical substrate, eIF2α. We could not use a *sui2-S51A* mutant strain that lacks the eIF2α Ser51 phosphorylation site since strains lacking the essential *SUI2* gene that encodes eIF2α, complemented with plasmids that express *SUI2* or *sui2-S51A* display constitutive PB formation even in the absence of stress (Supplementary Fig. 1C). Presumably, an imbalance in this essential translation initiation factor means that cells are constitutively stressed resulting in PB formation. Nevertheless, we did observe an oxidative stress-induced increase in PB formation in the *sui2-S51A* confirming that PB formation can occur independently of eIF2-phosphorylation. We therefore examined whether other components of the ISR are required for PB formation. Gcn1 and Gcn20 are required for Gcn2 activation during amino acid starvation and oxidative stress conditions (Marton et al., 1997, Shenton et al., 2006). We found that like the *gcn2* mutant, loss of *GCN1* or *GCN20* either partially or completely prevented PB formation during oxidative stress and amino acid starvation conditions respectively (Fig. 3A and B). Gcn1 and Gcn20 were also not required for PB formation during glucose starvation conditions. Immunoblot analysis confirmed that loss of *GCN1* abolished eIF2α phosphorylation, whereas loss of *GCN20* diminished it (Fig. 3C), consistent with previous reports (Marton et al., 1993, Vazquez de Aldana et al., 1995).

**Figure 3.**
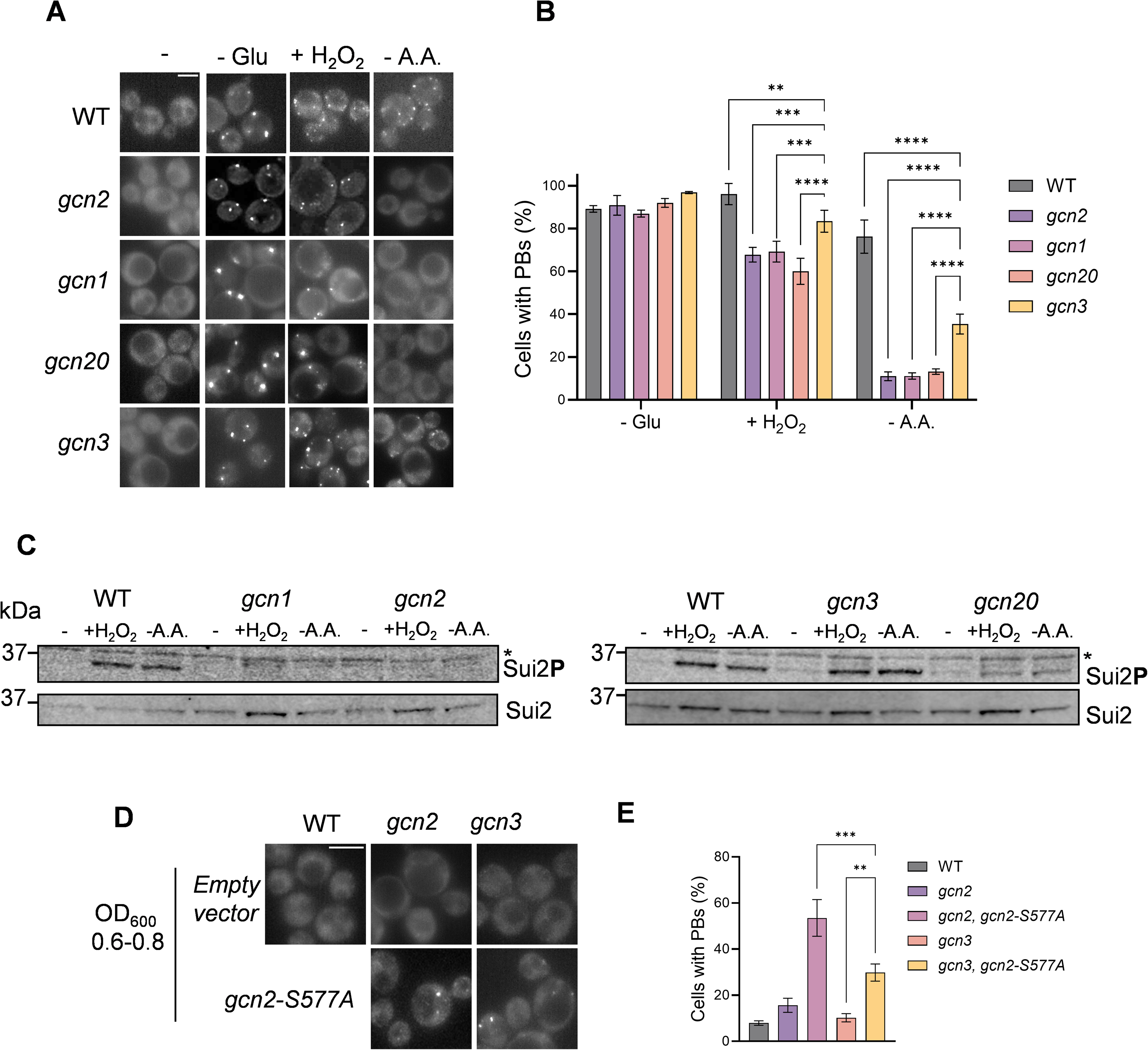
Requirement for components of the ISR in PB formation. **A.** Representative epifluorescent microscopic images are shown for wild-type, *gcn2, gcn1, gcn20* and *gcn3* mutant strains expressing Dcp1-GFP. Strains were grown to exponential phase and left untreated (-), starved for glucose (-Glu), starved for ammino acids (-AA) or treated with hydrogen peroxide (+H_2_O_2_). Scale bar 4 µm. **B.** Charts show the percentage of cells containing Dcp1-GFP foci (*n* > 100, based on three biological replicates). Error bars denote standard deviation. Significance is shown using a one-way ANOVA test; ** < 0.01, *** < 0.005, **** < 0.001. **C.** Phosphorylation of eIF2α is shown for strains grown as above. Western blots are shown probed with antibodies that recognise eIF2α-P (Sui2P) or eIF2α (Sui2) as a loading control. Asterisks denote a non-specific band recognized by anti-eIF2α-P. **D.** Representative epifluorescent microscopic images are shown for the wild-type strain containing vector (WT) and the *gcn2* and *gcn3* mutants containing vector or vectors expressing *gcn2-S577A.* **E.** Charts show the percentage of cells containing Dcp1-GFP foci (*n* > 100, based on three biological replicates). Error bars denote standard deviation. Significance is shown using a one-way ANOVA test; ** < 0.01, *** < 0.005.

To inhibit translation initiation, the phosphorylated form of eIF2α binds tightly to eIF2B, the guanine nucleotide exchange factor required for the recycling of eIF2, effectively sequestering eIF2B and diminishing its availability to regenerate eIF2-GTP (Kimball et al., 1998, Krishnamoorthy et al., 2001). Gcn3 is the alpha subunit of the eIF2B complex and is required to modulate the inhibition of eIF2B by eIF2α-P. We therefore examined PB formation in a *gcn3* mutant strain to test the reliance on eIF2α-P formation (Fig. 2A). Loss of *GCN3* was found to partially abrogate the induction of PB formation observed in response to oxidative stress and amino acid starvation but not to the same extent as the loss of *GCN2* (Fig. 3B). Immunoblot analysis confirmed that loss of *GCN3* does not affect eIF2 phosphorylation during these stress conditions (Fig. 3C). These data suggest that Gcn2 may be able to promote PB formation independently of its role in phosphorylating eIF2α. To further test the idea, PB formation was examined in a *gcn3* mutant strain expressing constitutively active *gcn2-S577A* (Fig. 3D and E). PB formation was significantly induced in this strain despite the absence of Gcn3 which is required for the binding eIF2α-P to eIF2B. Taken together these data suggest that Gcn2 can partially induce PB formation independently of its function in phosphorylating eIF2α.

### Identification of potential Gcn2-substrate proteins required for PB formation

A recent quantitative phosphoproteomics study identified 15 potential Gcn2 phospho-targets in addition to eIF2α (Dokladal et al., 2021). Based on this study, we used a candidate approach to identify Gcn2-substrate proteins required for PB formation. We focussed on non-essential proteins that have potential connections with protein synthesis including Sap185, Ssd1, Ubp3, Yar1, and Smu2. Interestingly, loss of each of these potential Gcn2-substrate proteins reduced PB formation induced by amino acid starvation, and loss of Ubp3 and Smu2 also reduced oxidative stress-induced PB formation (Fig. 4). Ubp3 is a ubiquitin-specific protease that has a well-characterized role in stress granule formation (Nostramo et al., 2016, Zhu et al., 2020). The only reported function for Smu2 is in targeting defective ribosomes for degradation by increasing the efficiency of the Fap1 E3 Ub-ligase which acts as a sensor of stalled ribosomes (Li et al., 2022). For the rest of this study, we focussed on the potential role of Smu2 in promoting PB formation.

**Figure 4.**
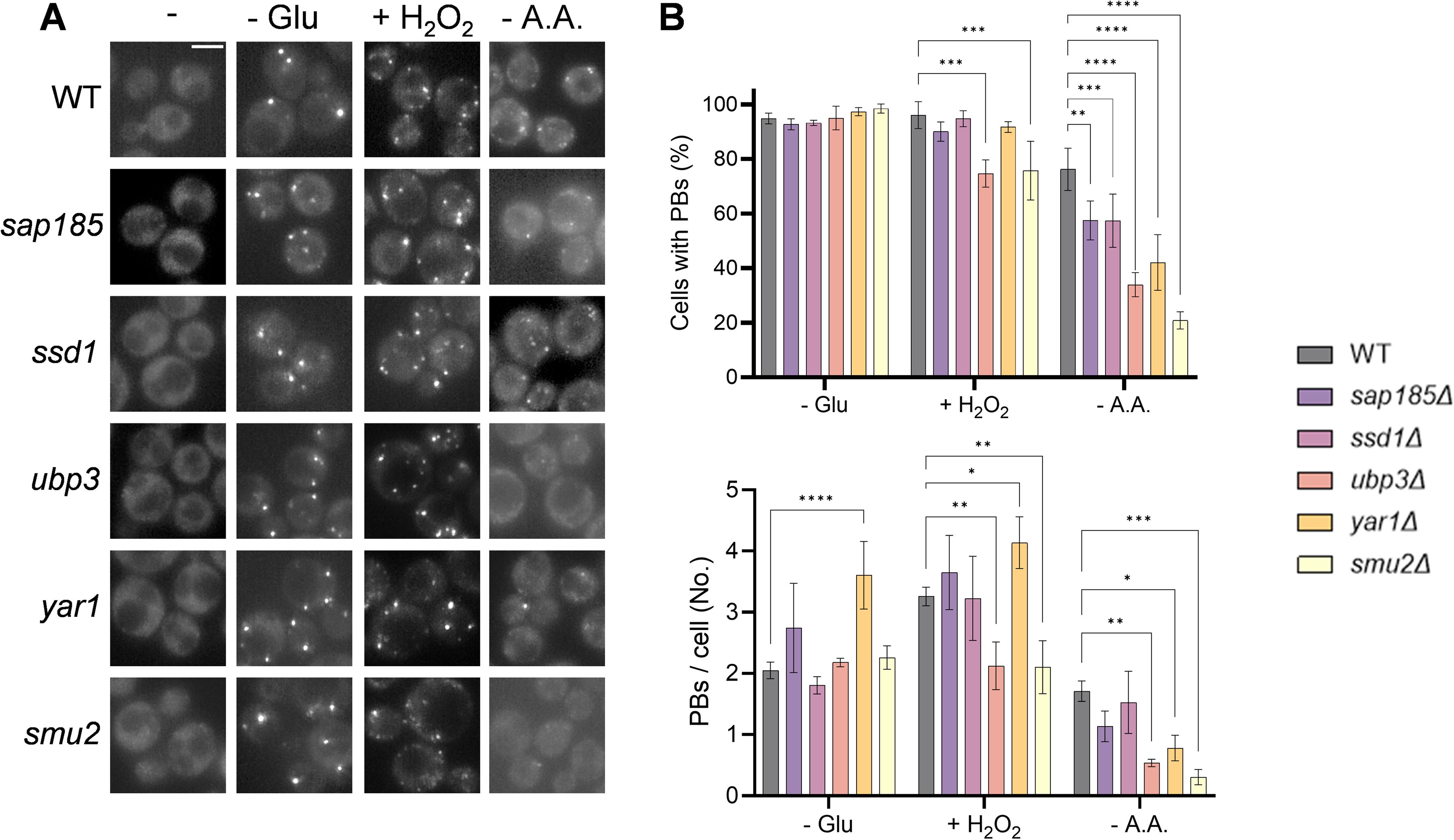
Identification of potential Gcn2-substrate proteins required for PB formation. **A.** Representative epifluorescent microscopic images are shown for wild-type, *sap185, ssd1, ubp3, yar1* and *smu2* mutant strains expressing Dcp1-GFP. Strains were grown to exponential phase and left untreated (-), starved for ammino acids (-AA) or treated with hydrogen peroxide (+H_2_O_2_). Scale bar 4 µm. **B.** Charts show the percentage of cells containing Dcp1-GFP foci and the average number of Dcp1-GFP foci per cell (*n* > 100, based on three biological replicates). Error bars denote standard deviation. Significance is shown using a one-way ANOVA test; * < 0.05, ** < 0.01, *** < 0.005, **** < 0.001.

### Smu2 is required for PB-formation independently of its function in uS3 ubiquitination

Ribosome stalling during translation can cause the production of truncated or dysfunctional proteins. To prevent this, ribosomal stalling triggers quality control responses, including ribosome ubiquitination. Smu2 and its co-interactors Fap1 and Fpr1 have been shown to trigger uS3 poly-ubiquitination in response to stalled 80S ribosomes that are initially mono-ubiquitinated by Mag2 (Li et al., 2022). Smu2 is thought to act as a cofactor for the Fap1 E3 ubiquitin ligase, which along with Fpr1, increases the efficiency of uS3 polyubiquitination on stalled monosomes (Fig. 5A).

**Figure 5.**
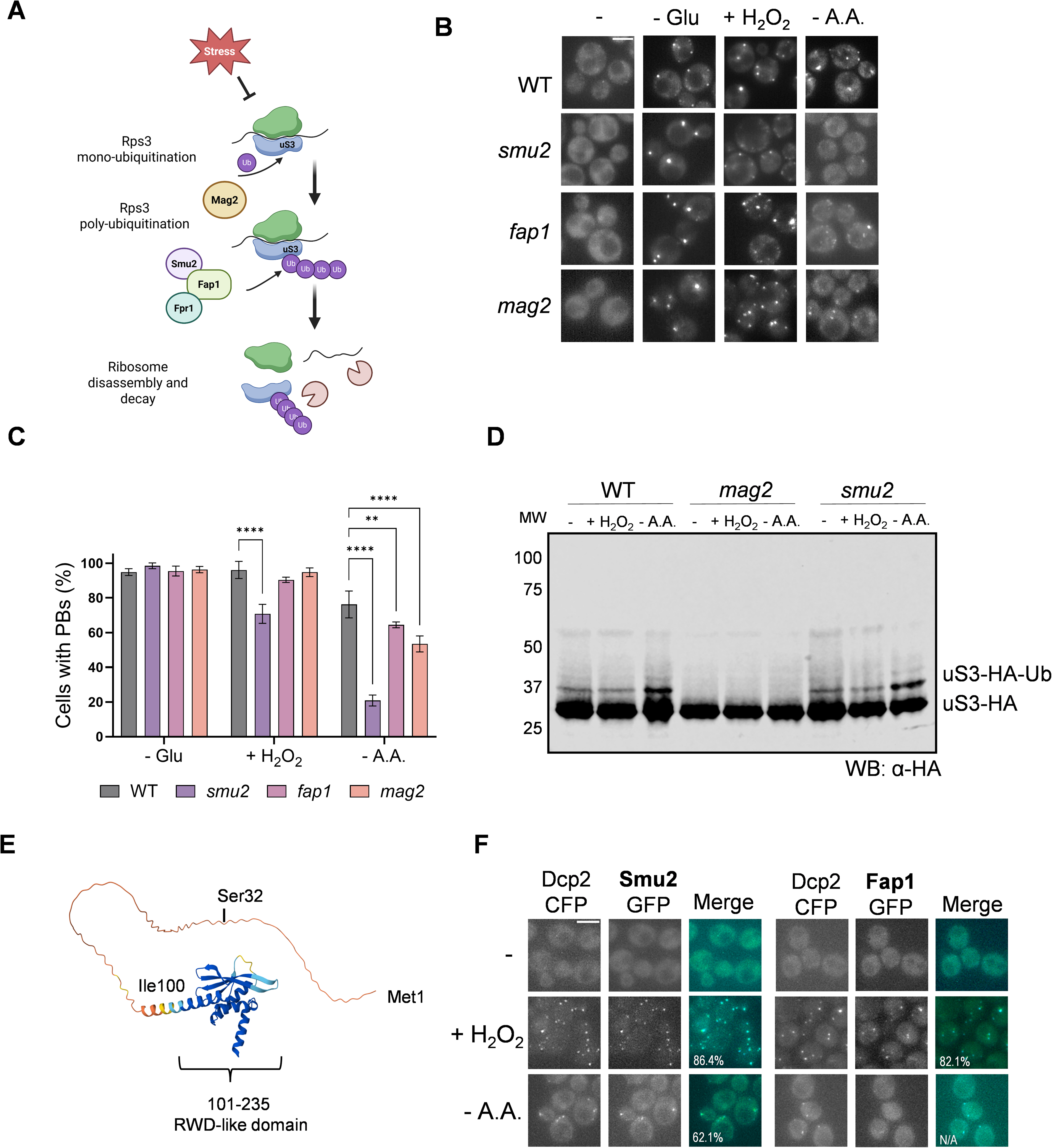
Smu2 is not required for PB formation via its function in Fap1-mediated uS3 ubiquitination. **A.** Smu2 functions as part of the Fap1 complex that recognises non-collided, but stalled 80S ribosomes (Li et al., 2022). Smu2 was identified as a co-factor that binds to stalled translating ribosomes along with the Fap1 Ub-ligase as part of the 18S nonfunctional rRNA decay (NRD) degradation pathway. Smu2 is not essential for uS3 polyubiquitination but was found to enhance its efficiency following mono ubiquitination by Mag2. **B.** Representative epifluorescent microscopic images are shown for wild-type, *smu2, fap1* and *mag2* mutant strains expressing Dcp1-GFP. Strains were grown to exponential phase and left untreated (-), starved for ammino acids (-AA) or treated with hydrogen peroxide (+H_2_O_2_). Scale bar 4 µm. **C.** Charts show the percentage of cells containing Dcp1-GFP foci (*n* > 100, based on three biological replicates). Error bars denote standard deviation. Significance is shown using a one-way ANOVA test; ** < 0.01, **** < 0.001. **D.** Whole cell extracts were prepared from wild-type, *mag2* and *smu2* mutant strains expressing uS3-HA. Western blots are shown probed with antibodies that recognise HA and the formation of uS3-HA-Ub is indicated. **E.** AlphaFold3 prediction of Smu2 structure showing a long unstructured N-terminal region (1–100) along with a compact C-terminal structure that contains a WD domain. Ser32 has been identified as the site of Gcn2 phosphorylation. **F.** Representative epifluorescent microscopic images are shown for strains expressing Dcp2-CFP and Smu2-GFP or Fap1.GFP. Numbers indicate the percentage of all Dcp1 or Smu2/Fap1 foci co-localised with each other (*n* > 100, based on three biological replicates).

We examined PB formation in mutants lacking the Fap1 E3 Ub ligase and found that PB formation was unaffected during oxidative stress conditions and only very moderately decreased during amino acid starvation conditions (Fig. 5B and C). Given this modest decrease, it seems unlikely that the mechanism by which Smu2 promotes PB formation involves Fap1. To assess this further, we examined uS3 ubiquitination via western blotting. We were unable to detect any alterations in the profile of uS3 in response to oxidative stress or amino acid starvation stress conditions (Fig. 5D). Mag2-dependent mono-ubiquitination of uS3 was detected but this too was unaffected by oxidative stress or amino acid starvation stress conditions (Fig. 4D). Probably the most telling result is the high levels of PB formation quantified in the *mag2* mutant, where no mono-ubiquitination of uS3 is detected (Fig. 5B and C). This supports the view that the function of Smu2 in ubiquitination is not required for PB formation.

Structural analysis of the Smu2 protein revealed that it contains a long N-terminal unstructured region (Li et al., 2022) (Fig. 5E). Given that regions of low sequence complexity have been implicated in the formation of stress-induced RNP granules, we examined whether Smu2 localizes to PBs (Youn et al., 2019). We found strong overlaps between Smu2-GFP and Dcp2-CFP granules formed in response to both oxidative stress (86%) and amino acid starvation conditions (64%) (Fig. 5F). In comparison, Fap1-GFP displayed high PB localisation in response to oxidative stress (82%), but no co-localisation was observed in response to amino acid starvation. Taken together, these data suggest that Smu2 is important for PB formation and its role is largely independent of Fap1 and the ubiquitination of uS3.

### Smu2 is required for mRNA localisation to RNP granules

One possibility to explain the requirement for Smu2 in stress-induced PB formation, is that it might function in the condensation of mRNAs to RNP granules. Given the links between translational repression and PB formation, we first tested whether loss of Smu2 has any effects on translational activity during stress conditions. Polysome analysis confirmed that loss of Smu2 does not affect the normal inhibition of translation initiation that is observed in response to amino acid starvation of oxidative stress conditions (Fig. 6A). Thus, the reduced PB formation that occurs in Smu2 mutants does not appear to arise due to any defects in the translational inhibition that promotes RNP granule formation. Additionally, loss of *SMU2* did not affect the phosphorylation of eIF2α that normally occurs in response to amino acid starvation and oxidative stress conditions (Fig. 6B).

**Figure 6.**
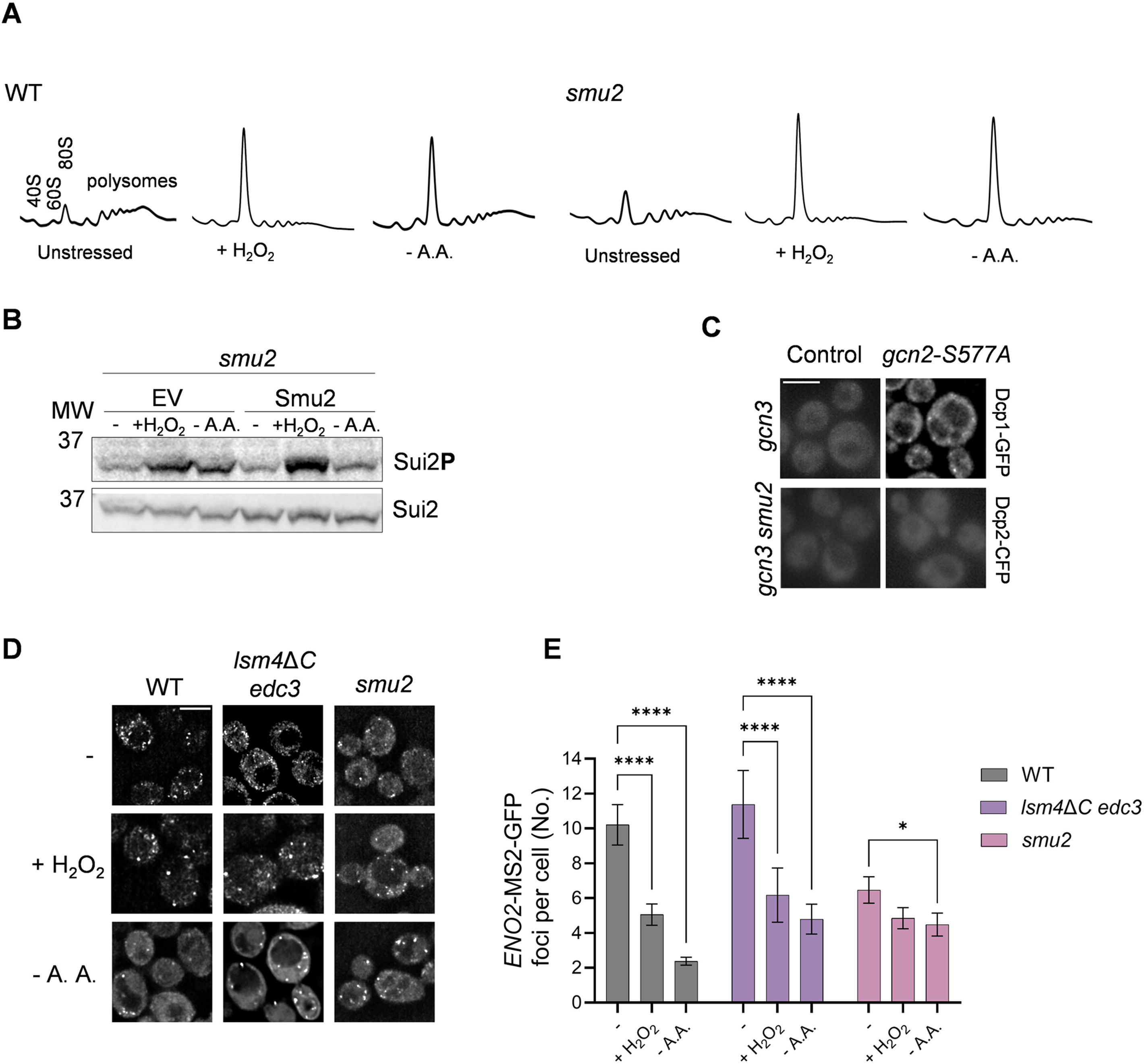
Smu2 is required for mRNA localisation to RNP granules. **A.** Representative polyribosome traces are shown for the wild-type and *smu2* mutant strains grown to exponential phase and left untreated, starved for ammino acids (-AA) or treated with hydrogen peroxide (+H_2_O_2_). The peaks that contain the small ribosomal subunit (40S), the large ribosomal subunit (60S), and both subunits (80S) are indicated along with the polysome peaks generated by 2, 3, 4, 5, etc., 80S ribosomes on a single mRNA. **B.** Phosphorylation of eIF2α is shown for the *smu2* mutant containing vector (EV) or containing a vector expressing Smu2. Western blots are shown probed with antibodies that recognise eIF2α-P (Sui2P) or eIF2α (Sui2) as a loading control. **C.** Representative epifluorescent microscopic images are shown for *gcn3* and *gcn3 smu2* strains expressing Dcp1-GFP and containing vector (Control) or vector expressing *gcn2-S577A.* Scale bar = 4 μm. **D.** *ENO2*-MS2 mRNA was visualised in wild-type, *lsm4*Δ*C edc3*Δ and *smu2* cells expressing GFP-fused coat protein. **E.** Charts show the average number of *ENO2-MS2-GFP* foci per cell (*n* > 100, based on three biological replicates). Error bars denote standard deviation. Significance is shown using a one-way ANOVA test; * < 0.05, **** < 0.001.

To further confirm, that Smu2 controls PB formation independently of eIF2α phosphorylation, PB formation was examined in a *gcn3 smu2* double mutant. Gcn3 enables the inhibition of eIF2B by eIF2α-P and loss of *GCN3* was found to partially abrogate the induction of PB formation caused by the constitutive *gcn2-S577A* mutant (Fig. 3A). Simultaneous loss of Gcn3 and Smu2 further reduced PB formation in the constitutive *gcn2-S577A* mutant (Fig. 6C). This additive effect confirms that Smu2 and eIF2α phosphorylation mediate PB formation independently.

To examine mRNA condensation in a *smu2* mutant, we again used the *ENO2* mRNA as a well characterized mRNA that localises to PBs (Lui et al., 2014, Pizzinga et al., 2019). For these experiments PB formation was also compared with the *edc3*Δ *lsm4*Δ*C* mutant which is defective in PB formation induced by amino acid starvation, oxidative stress and glucose depletion (Fig. 1D). Interestingly, the number of *ENO2* mRNA granules observed under non-stress conditions was decreased in the *smu2* mutant compared with both the wild-type and *lsm4*Δ*C edc3* mutant strains (Fig. 7D). The normal reduction in the numbers of *ENO2* mRNA granules that occurs in the wild-type and *lsm4*Δ*C edc3* mutant strains when mRNAs condense into PBs was therefore more muted in response to amino acid starvation and abrogated in response to oxidative stress in *smu2* mutant strains. Thus, although translation inhibition still occurs in strains lacking *SMU2*, it appears that less mRNA material is sequestered into PBs following stress conditions, which may explain the reduced frequency of PB formation in response to amino acid starvation and oxidative stress.

**Figure 7.**
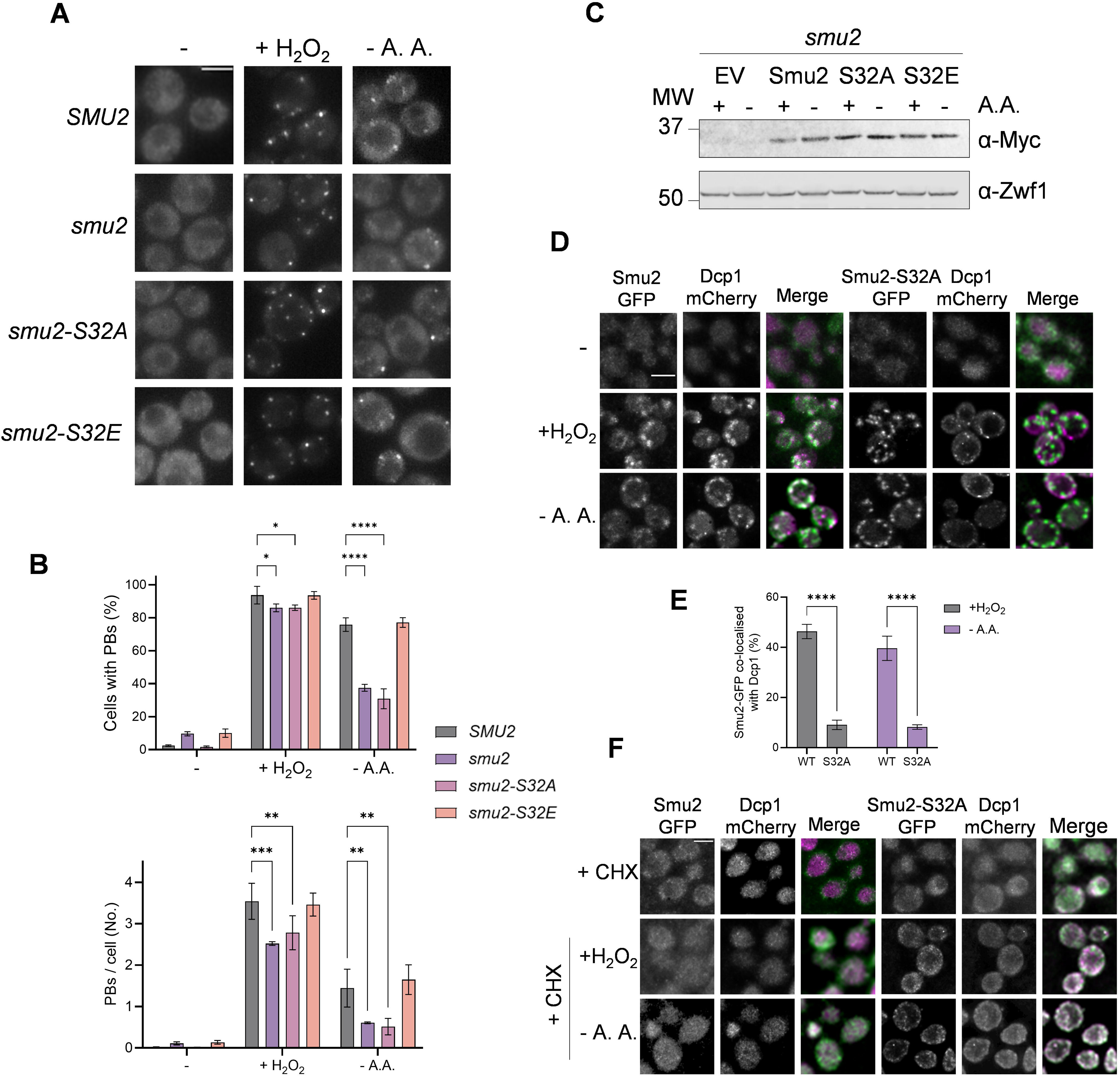
The Ser32 Gcn2-phosphorylation site in Smu2 is required for Smu2 localisation and PB formation. A. Representative epifluorescent microscopic images are shown for *smu2* mutant strains expressing Dcp1-GFP and containing vector (*smu2*) or vector expressing *SMU2, smu2-S32A or smu2-S32E*. Strains were grown to exponential phase and left untreated (-), starved for ammino acids (-AA) or treated with hydrogen peroxide (+H_2_O_2_). Scale bar 4 µm. **B.** Charts show the percentage of cells containing Dcp1-GFP foci and the average number of Dcp1-GFP foci per cell (*n* > 100, based on three biological replicates). Error bars denote standard deviation. Significance is shown using a one-way ANOVA test; * < 0.05, ** < 0.01, *** < 0.005, **** < 0.001. **C.** Smu2 protein levels are shown for the *smu2* mutant containing vector (EV) or containing a vector expressing Smu2, Smu2-S32A or Smu2-S32E. Western blots are shown probed with antibodies that recognise Smu2-Myc or Zwf1 as a loading control. **D.** Representative epifluorescent microscopic images are shown for *smu2* mutant strains expressing Dcp1-mCherry and containing vector expressing Smu2-GFP or Smu2-S32A-GFP. Dcp1-mCherry is shown in magenta. **E.** Charts show the percentage of Smu2-GFP foci that co-localise with Dcp1-mCherry foci (*n* > 100, based on three biological replicates). Error bars denote standard deviation. Significance is shown using an unpaired t-test, **** < 0.001. **F.** Same as for (E) except cells were treated with cycloheximide (10 min, 0.1 mg/ml) prior to imposing stress conditions.

### The Gcn2-phosphorylation site in Smu2 is required for PB formation

Previous phosphoproteomic studies identified Ser32 in Smu2 as the site of Gcn2-mediated phosphorylation (Dokladal et al., 2021). This phosphorylation site is in the long N-terminal unstructured region of Smu2, and hence phosphorylation may affect Smu2 PB localisation (Fig. 5E). To test this possibility, plasmids expressing *SMU2,* a non-phosphorylatable *smu2-S32A* mutant and a phosphomimetic *smu2-S32E* mutant were constructed and introduced into a *smu2* mutant strain. The *smu2* deletion strain expressing Smu2-S32A showed reduced numbers of cells with PBs and reduced numbers of PBs per cell during oxidative stress and amino acid starvation conditions comparable to the deletion strain (Fig. 7A and B). The strain expressing the phosphomimetic *smu2-S32E* mutant exhibited a similar pattern of PB formation to the strain expressing wild-type *SMU2*. The *smu2-S32E* mutant did not form PBs in the absence of stress, suggesting that although the phosphorylation site is required for stress induced PB formation, a phosphomimetic mutant is not sufficient to promote PB formation (Fig. 7A and B). These data indicate that Ser32 in Smu2 is important for stress-induced PB formation. We confirmed that that the Smu2, Smu2-S32A and Smu2-S32E mutants are expressed at comparable levels using immunoblot analysis (Fig. 7C). Additionally, no changes in Smu2 expression were observed following amino acid starvation conditions.

Post-translation modifications are important for many regulated protein localisation events within cells, and more specifically can affect protein recruitment or exclusion from RNP granules (Lee et al., 2023, Wang et al., 2023). Having established that a non-phosphorylatable *smu2-S32A* mutant suppresses PB formation, we next tested whether Ser32 is important for Smu2 localisation to PBs. To test this, C-terminal GFP fusion constructs were generated for wild-type Smu2 and the non-phosphorylatable smu2-S32A mutant. As shown previously (Fig. 6F), wild-type Smu2 was again found to localise to PBs in response oxidative stress and amino acid starvation conditions (Fig. 7D and E). Interestingly, the non-phosphorylatable Smu2-S32A protein still formed multiple granules in response to oxidative stress and amino acid starvation, but co-localisation with PBs was strongly reduced suggesting that the proportion of Smu2 in non-PB granules increases for the Smu2-S32A mutant protein (Fig. 7D and E). The identity of the non-PB Smu2 granules is unclear, but our initial experiments found that treating cells with the protein synthesis inhibitor cycloheximide prevented the localisation of Smu2-S32A to both PBs and these novel granules. Inhibiting translation elongation using cycloheximide is known to prevent RNP granule formation by preventing the release of mRNAs from polysomes (Teixeira et al., 2005, Buchan et al., 2008), suggesting that these granules do not arise due to misfolding and aggregation of mutant Smu2-S32A (Fig. 7F). Taken together, these data are consistent with a role for Ser32 phosphorylation in Smu2 localisation to PBs which facilitates PB formation.

## Discussion

PBs are cytoplasmic ribonucleoprotein granules that function as dynamic sites of mRNA storage, translational repression and degradation (Jain and Parker, 2013, Luo et al., 2018, Standart and Weil, 2018). They form when translation initiation is inhibited and non-translating mRNAs accumulate outside the actively translated pool (Teixeira et al., 2005, Teixeira and Parker, 2007). During stress, this reversible sequestration protects and stores mRNAs until they can leave PBs and re-engage with the translational machinery, thereby coordinating mRNA metabolism with protein synthesis (Brengues et al., 2005, Horvathova et al., 2017, Feng et al., 2026). Here, we show that amino acid starvation, oxidative stress and glucose starvation induce PBs with similar marker localisation, mRNA sequestration and biophysical properties. However, the upstream signalling requirements differ: glucose-starvation-induced PB formation is independent of Gcn2, amino acid starvation-induced PB formation strongly depends on Gcn2, and oxidative-stress-induced PB formation depends only partially on Gcn2.

Inhibition of translation initiation is a conserved response to diverse stresses. In yeast, Gcn2 is the sole eIF2α kinase and inhibits translation initiation through eIF2α phosphorylation during oxidative stress and amino acid starvation (Dever et al., 1992, Shenton et al., 2006). By contrast, glucose starvation rapidly inhibits translation initiation independently of eIF2α phosphorylation by disrupting the interaction between eIF4A and the 48S preinitiation complex, explaining why glucose-starvation-induced PB formation does not require Gcn2 (Ashe et al., 2000, Castelli et al., 2011, Bresson et al., 2020). The different requirements for Gcn2 during amino acid starvation and oxidative stress are less readily explained. Although Gcn2 is required for most of the inhibition of translation initiation under both conditions, its loss abolishes PB formation during amino acid starvation but only partially reduces it during oxidative stress. Oxidative damage to mRNAs and the translation machinery may provide an additional route to PB formation. Oxidative stress inhibits translation elongation and promotes the accumulation of stalled and collided ribosomes on mRNAs (Shenton et al., 2006, Simms et al., 2014, Wu et al., 2019). These stalled ribosomes activate a conserved ribosome-associated quality-control pathway that ultimately targets damaged or stalled mRNAs for degradation (Inada, 2017, D’Orazio and Green, 2021, Yip and Shao, 2021, Kim and Zaher, 2022). Damaged mRNAs may therefore promote PB condensation if they accumulate or contain ribosome-free regions downstream of stall sites. Consistent with this possibility, oxidised RNAs localise to cytoplasmic condensates implicated in translation quality control that also contain several PB and stress-granule markers (Dhaliwal et al., 2022). Because mRNA recruitment to PBs depends on both physical properties and the inducing stress (Khong et al., 2017, Kershaw et al., 2021), PBs formed during oxidative stress and amino acid starvation are likely to contain different mRNA repertoires and perform stress-specific functions.

Expression of a constitutively active Gcn2 allele induces PB formation in the absence of exogenous stress (Buchan et al., 2008). Surprisingly, we found that these PBs are still formed in a *gcn3* mutant, in which eIF2α phosphorylation does not inhibit eIF2B, suggesting that Gcn2 might promote PB formation through additional, eIF2α-independent mechanisms. Similarly, mammalian stress granules can form independently of eIF2α phosphorylation in a *sui2-S52A* mutant during hydrogen peroxide stress (Emara et al., 2012). Recent studies have identified additional Gcn2 substrates, supporting functions beyond activation of the integrated stress response (ISR) (Ge et al., 2023, Stonyte et al., 2023). Phosphoproteomic analysis identified 16 putative direct Gcn2 substrates, including Gcn20 and Sui3, the β-subunit of eIF2, which regulate translation initiation (Dokladal et al., 2021). Other putative substrates include metabolic enzymes and proteins with established roles in RNA metabolism and translational control. One example is Pat1, an RNA-binding protein that promotes PB formation (Sachdev et al., 2019). These candidate substrates provide plausible routes through which Gcn2 could regulate PB formation independently of eIF2α phosphorylation.

Using a candidate approach, we examined PB formation in mutants lacking the putative Gcn2 substrates Sap185, Ssd1, Ubp3, Yar1 and Smu2. Loss of each protein reduced PB formation to some extent, although the mechanisms may differ. Ubp3 is a ubiquitin-specific protease with a well-characterised role in stress-granule formation and may therefore have a broader function in RNP-granule assembly, including PB formation (Nostramo et al., 2016, Zhu et al., 2020). Sap185 associates with the Sit4 type 2A-related protein phosphatase and has been implicated in eIF2α dephosphorylation, suggesting that it could affect PB formation by altering eIF2α phosphorylation (Luke et al., 1996, Rohde et al., 2004). Ssd1 is an mRNA-binding protein involved in transcript-specific translation and age-dependent condensate formation (Luukkonen and Seraphin, 1999, Jansen et al., 2009, Gutierrez et al., 2026), whereas Yar1 is a dedicated chaperone for the ribosomal protein Rps3 (Koch et al., 2012). However, a possible explanation for the influence of Ssd1 and Yar1 on PB formation remains unclear. Together, these findings extend the links between Gcn2-regulated proteins, RNA metabolism and RNP-granule formation.

We focused on Smu2 since it has the most pronounced defect in PB formation. Smu2 is a component of the Fap1 complex that recognises non-collided but stalled 80S ribosomes (Li et al., 2022). In the 18S nonfunctional rRNA decay pathway, Smu2 binds stalled translating ribosomes with the Fap1 ubiquitin ligase and enhances uS3 polyubiquitination after initial monoubiquitination by Mag2. However, we detected no uS3 polyubiquitination following oxidative stress or amino acid starvation, and loss of Mag2 or Fap1 had only modest effects on PB formation. These findings indicate that Smu2 promotes PB formation independently of its established role in uS3 ubiquitination and nonfunctional rRNA decay. Loss of Smu2 suppressed PB formation to a similar extent as loss of Gcn3, while combined loss of Gcn3 and Smu2 caused a further reduction. This additive effect indicates that Smu2 and the eIF2α-dependent pathway act independently downstream of Gcn2 to promote PB formation.

Smu2 contains a long intrinsically disordered N-terminal region spanning residues 1– 100 and a compact C-terminal region containing a WD domain. WD domains are often found in RING finger-containing proteins, WD repeat-containing proteins and DEAD (DEXD)-like helicases (Li et al., 2022). Intrinsically disordered regions (IDRs) frequently contribute to the localisation and assembly of stress-induced RNP granules (Protter et al., 2018, Youn et al., 2019). Consistent with this property, Smu2 localised to the PBs formed during both oxidative stress and amino acid starvation. Ser32, the reported site of Gcn2-mediated phosphorylation, lies within the N-terminal IDR (Dokladal et al., 2021; Fig. 5E). Because phosphorylation within IDRs can alter their structure and interactions with proteins and RNAs, Ser32 phosphorylation could regulate Smu2 recruitment to PBs (Modic et al., 2024). Indeed, mutation of Ser32 suppressed PB formation under both stresses to a similar extent as complete loss of Smu2, and the non-phosphorylatable S32A protein showed reduced localisation to PBs. Proteomic screens have also identified Smu2 as an RNA-binding protein (Beckmann et al., 2015), consistent with the enrichment of RNA-binding domains and IDRs among RNP-granule proteins (Sanders et al., 2020, Yang et al., 2020, Kershaw et al., 2021). Together, these findings support a model in which Gcn2-dependent phosphorylation of Ser32 promotes Smu2 interactions with RNA or other RNP components, facilitating its recruitment to PBs and contributing to condensate formation.

Translationally repressed mRNAs localise to RNP granules such as PBs and stress granules during stress, but translated mRNAs also form granules in unstressed cells (Lui et al., 2014, Pizzinga et al., 2019, Morales-Polanco et al., 2021). These condensates include translation factories in actively growing cells. For example, *ENO2* mRNA, which encodes enolase, is normally present in more than 10 Core Fermentation granules per cell (Morales-Polanco et al., 2021). During stress, pre-existing translation factories are remodelled and coalesce with other proteins and RNAs to seed PB formation (Lui et al., 2014). Consistent with this model, we observed *ENO2* mRNA granules coalescing into fewer, larger granules during oxidative stress and amino acid starvation, coincident with recruitment of PB components. This coalescence persisted in the PB-assembly-defective *lsm4*Δ*C edc3*Δ mutant, as previously reported during glucose starvation (Lui et al., 2014), indicating that mRNA condensation can precede full PB assembly. By contrast, loss of Smu2 reduced the number of *ENO2* mRNA granules in unstressed cells and largely abolished the stress-induced change in granule number. Smu2 may therefore maintain the pre-existing mRNA condensates that seed PB formation and promote recruitment of PB components during stress. Overall, our findings support two complementary routes through which Gcn2 promotes PB formation: eIF2α phosphorylation increases the pool of untranslated mRNAs, while Smu2 promotes the maturation of pre-existing mRNA granules into PBs.

## Materials and Methods

### Yeast strains and plasmids

*Saccharomyces cerevisiae* strains used in this study are listed in Supplementary Table 1 and were all derivatives of W303-1A (*MATa ura3-52 leu2-3 leu2-112 trp1-1 ade2-1 his3-11 can1-100*). Gene deletion and C-terminal tagged strains were constructed using standard yeast methodology (Carter and Delneri, 2010, Knop et al., 1999) Plasmids containing *GCN2*, *gcn2-m2*, and *gcn2-S577A* have been described previously (Cherkasova and Hinnebusch, 2003). Plasmids expressing Smu2-Myc, Smu2-S32A-Myc, Smu2-S32E-Myc, Smu2-GFP and Smu2-S32A-GFP were constructed by cloning commercially synthesized gene fragments into plasmid pRS414 (Sikorski and Hieter, 1989). The plasmid expressing Dcp1-mCherry has been described previously (Nostramo et al., 2019).

### Growth and stress conditions

Yeast strains were grown at 30°C with shaking at 180 rpm in SCD complete media (0.67% w/v yeast nitrogen base without amino acids, 2% w/v glucose) supplemented with appropriate amino acids and bases (ForMedium, UK). All experiments were performed using cells grown to mid-log phase (OD_600_ 0.5-0.8). Amino acid and glucose starvation conditions were induced by resuspending cells in prewarmed media lacking all amino acids or lacking glucose. Oxidative stress was imposed by adding H_2_O_2_ to a final conc. of 0.8 mM. To visualise the effects of cycloheximide on foci formation, cells were pre-treated with cycloheximide (0.1 mg/ml) for 10 minutes prior to stress conditions. Cells were resuspended in media containing 5% (w/v) 1,6-hexanediol (Merck Life Science, UK) to dissolve condensates.

### Live cell imaging

Confocal images were taken using a LeicaDM550 B or a Nikon Eclipse Ni-E microscope. The Leica microscope was equipped with Leica K5 cMOS camera and Leica100x/1.40-0.70NA oil plan Apo objective, whereas the Nikon microscope was equipped with a DS-Qi2 mono digital camera and 100x/1.3 NA oil objective. The fluorescent parameters for both microscopes were: CFP excitation 436/20 nm, emission 480/40 nm, GFP excitation 470/40 nm, emission 525/50 nm, RFP excitation 560/40nm, emission 645/75 nm. Live yeast cells were plated on poly-L-lysine coated slides and images were taken with a z-spacing of 200 nm. Images were processed using ImageJ (Schneider et al., 2012) and quantification of the PBs was performed manually and using the ImageJ plugin ComDet (v.0.5.5; Katrukha, 2022). Raw images were deconvolved using the Huygens Pro software (SVI) and maximum intensity projections of these deconvolved images are shown in the results.

### Protein extraction and western blotting

Protein extracts were electrophoresed under reducing conditions on NuPAGE SDS-PAGE minigels (ThermoFisher Scientific) and electroblotted onto PVDF membrane (Amersham Pharmaica Biotech) using standard wet transfer. Primary antibodies used recognised Myc (Myc4A6, Millipore), Pgk1 (Abcam ab113687), HA (Roche 11867423001), Zwf1 (Sigma-Aldrich A9521), Sui2 (Biorbyt orb847328) and Sui2-P (Proteintech 68023).

### Polysome profiling and analysis

Polysome profiling was performed as previously described (Ashe et al., 2000). Briefly, cells were grown to OD_600_ of 0.6 before fixing ribosomes on mRNAs by treatment with 0.1mg/ml cycloheximide on ice-cold water for 10 minutes. Cells were lysed with acid-washed glass beads in lysis buffer (20 mM HEPES pH 7.4, 100 mM potassium acetate, 2 mM magnesium acetate, 0.5 mM DTT, 0.4 mM cycloheximide) by vortexing for 20 seconds, 8-10 times. 2.5 *A*_260_ units were layered onto 15-50% sucrose gradients prepared in 12 ml thin-walled open polyallomer tubes (Seton Scientific) and separated by ultracentrifugation in an SW41 Ti rotor (2.5 h at 278 000 × g, 4°C). Polysome profiles were generated using a TRIAX™ flow cell UV/Vis recorder continuously monitoring *A*_260_. Polysome traces and polysome to monosome ratios were generated using QuAPPro (v. 0.3.1).

## Acknowledgements

We thank Graham Pavitt and the University of Manchester Bioimaging facility for helpful advice and discussions. We thank Toshi Inada for the *uS3-3HA::hisMX6* strain and Alan Hinnebusch for the plasmids expressing *GCN2, gcn2-m2*, and *gcn2-S577A*.

## Footnotes

### AI Declaration

Copilot was used to improve human-generated texts for readability and style

### Competing interests

All strains and plasmids are available upon request. All relevant data and details of resources can be found within the article and its supplementary information.

### Funding

This work was supported by a Biotechnology and Biological Sciences Research Council (BBSRC) DTP studentship BB/T008725/1 to V.S.

### Data and resource availability

The authors declare no competing or financial interests.

### Author contributions

Conceptualization: C.M.G, M.P.A., S.J.H.; Funding acquisition: C.M.G, M.P.A., S.J.H.; Investigation: V.S.; Methodology: C.M.G, M.P.A., S.J.H., V.S.; Formal analysis: C.M.G, M.P.A., V.S.; Writing – original draft: C.M.G, V.S.; Writing – review and editing: C.M.G, M.P.A., S.J.H., V.S.

## Figure Legends

**Figure S1. Gcn2 is required for the inhibition of translation initiation in response to oxidative stress and amino acid starvation conditions.** Wild-type and *gcn2* mutant strains were grown to exponential phase and left untreated (-), starved for glucose (-Glu), starved for ammino acids (-AA) or treated with 0.8 mM hydrogen peroxide (+H_2_O_2_) all for one hour. **A.** Western blots are shown probed with antibodies that recognise eIF2α-P (Sui2P) or eIF2α (Sui2) as a loading control. **B.** Polyribosome traces are shown. The peaks that contain the small ribosomal subunit (40S), the large ribosomal subunit (60S), and both subunits (80S) are indicated along with the polysome peaks generated by 2, 3, 4, 5, etc., 80S ribosomes on a single mRNA. **C.** Dcp1-GFP was visualised in *SUI2* deleted strain complemented with plasmids expressing wild-type *SUI2* or a non-phosphorylatable *sui2-S51A* plasmids. Yeast cells visualised prior and post oxidative stress conditions.

## Notes

### Competing Interest Statement

The authors have declared no competing interest.

## References

Ashe, M. P., De Long, S. K. & Sachs, A. B. 2000. Glucose depletion rapidly inhibits translation initiation in yeast. Mol Biol Cell, 11, 833–48.

Bresson, S., Shchepachev, V., Spanos, C., Turowski, T. W., Rappsilber, J. & Tollervey, D. 2020. Stress-Induced Translation Inhibition through Rapid Displacement of Scanning Initiation Factors. Mol Cell, 80, 470–484 e8.

Buchan, J. R., Muhlrad, D. & Parker, R. 2008. P bodies promote stress granule assembly in Saccharomyces cerevisiae. J Cell Biol, 183, 441–55.

Carter, Z. & Delneri, D. 2010. New generation of loxP-mutated deletion cassettes for the genetic manipulation of yeast natural isolates. Yeast, 27, 765–75.

Cary, G. A., Vinh, D. B., May, P., Kuestner, R. & Dudley, A. M. 2015. Proteomic Analysis of Dhh1 Complexes Reveals a Role for Hsp40 Chaperone Ydj1 in Yeast P-Body Assembly. G3 (Bethesda), 5, 2497-511.

Castelli, L. M., Lui, J., Campbell, S. G., Rowe, W., Zeef, L. A., Holmes, L. E., Hoyle, N. P., Bone, J., Selley, J. N., Sims, P. F. & Ashe, M. P. 2011. Glucose depletion inhibits translation initiation via eIF4A loss and subsequent 48S preinitiation complex accumulation, while the pentose phosphate pathway is coordinately up-regulated. Mol Biol Cell, 22, 3379–93.

Castilho, B. A., Shanmugam, R., Silva, R. C., Ramesh, R., Himme, B. M. & Sattlegger, E. 2014. Keeping the eIF2 alpha kinase Gcn2 in check. Biochim Biophys Acta, 1843, 1948–68.

Chang, W. D. & Choe, Y. J. 2026. Quality control and signaling pathways at stalled ribosomes. Exp Mol Med, 58, 82–93.

Cherkasova, V. A. & Hinnebusch, A. G. 2003. Translational control by TOR and TAP42 through dephosphorylation of eIF2alpha kinase GCN2. Genes Dev, 17, 859–72.

Clemens, M. J. 2001. Initiation factor eIF2 alpha phosphorylation in stress responses and apoptosis. Prog Mol Subcell Biol, 27, 57–89.

Decker, C. J., Teixeira, D. & Parker, R. 2007. Edc3p and a glutamine/asparagine-rich domain of Lsm4p function in processing body assembly in Saccharomyces cerevisiae. J Cell Biol, 179, 437–49.

Dever, T. E., Feng, L., Wek, R. C., Cigan, A. M., Donahue, T. F. & Hinnebusch, A. G. 1992. Phosphorylation of initiation factor 2 alpha by protein kinase GCN2 mediates gene-specific translational control of GCN4 in yeast. Cell, 68, 585–96.

Dever, T. E., Kinzy, T. G. & Pavitt, G. D. 2016. Mechanism and Regulation of Protein Synthesis in Saccharomyces cerevisiae. Genetics, 203, 65–107.

Dokladal, L., Stumpe, M., Pillet, B., Hu, Z., Garcia Osuna, G. M., Kressler, D., Dengjel, J. & De Virgilio, C. 2021. Global phosphoproteomics pinpoints uncharted Gcn2-mediated mechanisms of translational control. Mol Cell, 81, 1879–1889 e6.

Donnelly, N., Gorman, A. M., Gupta, S. & Samali, A. 2013. The eIF2alpha kinases: their structures and functions. Cell Mol Life Sci, 70, 3493–511.

Feng, M. W., Mossanen-Parsi, A., Stonys, V., Hubbard, S. J., Ashe, M. P. & Grant, C. M. 2026. P-bodies act as dynamic control hubs for RNA processing and storage. J Biol Chem, 302, 111273.

Gupta, R. & Hinnebusch, A. G. 2023. Differential requirements for P stalk components in activating yeast protein kinase Gcn2 by stalled ribosomes during stress. Proc Natl Acad Sci U S A, 120, e2300521120.

Haim-Vilmovsky, L., Gadir, N., Herbst, R. H. & Gerst, J. E. 2011. A genomic integration method for the simultaneous visualization of endogenous mRNAs and their translation products in living yeast. RNA, 17, 2249–55.

Hinnebusch, A. G. 2005. Translational regulation of GCN4 and the general amino acid control of yeast. Annu Rev Microbiol, 59, 407–50.

Hinnebusch, A. G. & Fink, G. R. 1983. Positive regulation in the general amino acid control of Saccharomyces cerevisiae. Proc Natl Acad Sci U S A, 80, 5374–8.

Hoyle, N. P., Castelli, L. M., Campbell, S. G., Holmes, L. E. & Ashe, M. P. 2007. Stress-dependent relocalization of translationally primed mRNPs to cytoplasmic granules that are kinetically and spatially distinct from P-bodies. J Cell Biol, 179, 65–74.

Iurlaro, R., Puschel, F., Leon-Annicchiarico, C. L., O’connor, H., Martin, S. J., Palou-Gramon, D., Lucendo, E. & Munoz-Pinedo, C. 2017. Glucose Deprivation Induces ATF4-Mediated Apoptosis through TRAIL Death Receptors. Mol Cell Biol, 37.

Jain, S. & Parker, R. 2013. The discovery and analysis of P Bodies. Adv Exp Med Biol, 768, 23–43.

Kedersha, N., Stoecklin, G., Ayodele, M., Yacono, P., Lykke-Andersen, J., Fritzler, M. J., Scheuner, D., Kaufman, R. J., Golan, D. E. & Anderson, P. 2005. Stress granules and processing bodies are dynamically linked sites of mRNP remodeling. J Cell Biol, 169, 871–84.

Kershaw, C. J., Nelson, M. G., Lui, J., Bates, C. P., Jennings, M. D., Hubbard, S. J., Ashe, M. P. & Grant, C. M. 2021. Integrated multi-omics reveals common properties underlying stress granule and P-body formation. RNA Biol, 18, 655–673.

Kimball, S. R., Fabian, J. R., Pavitt, G. D., Hinnebusch, A. G. & Jefferson, L. S. 1998. Regulation of guanine nucleotide exchange through phosphorylation of eukaryotic initiation factor eIF2alpha. Role of the alpha- and delta-subunits of eiF2b. J Biol Chem, 273, 12841–5.

Knop, M., Siegers, K., Pereira, G., Zachariae, W., Winsor, B., Nasmyth, K. & Schiebel, E. 1999. Epitope tagging of yeast genes using a PCR-based strategy: more tags and improved practical routines. Yeast, 15, 963–72.

Krishnamoorthy, T., Pavitt, G. D., Zhang, F., Dever, T. E. & Hinnebusch, A. G. 2001. Tight binding of the phosphorylated alpha subunit of initiation factor 2 (eIF2alpha) to the regulatory subunits of guanine nucleotide exchange factor eIF2B is required for inhibition of translation initiation. Mol Cell Biol, 21, 5018–30.

Kroschwald, S., Maharana, S., Mateju, D., Malinovska, L., Nuske, E., Poser, I., Richter, D. & Alberti, S. 2015. Promiscuous interactions and protein disaggregases determine the material state of stress-inducible RNP granules. Elife, 4, e06807.

Lageix, S., Zhang, J., Rothenburg, S. & Hinnebusch, A. G. 2015. Interaction between the tRNA-binding and C-terminal domains of Yeast Gcn2 regulates kinase activity in vivo. PLoS Genet, 11, e1004991.

Lee, J. M., Hammaren, H. M., Savitski, M. M. & Baek, S. H. 2023. Control of protein stability by post-translational modifications. Nat Commun, 14, 201.

Li, S., Ikeuchi, K., Kato, M., Buschauer, R., Sugiyama, T., Adachi, S., Kusano, H., Natsume, T., Berninghausen, O., Matsuo, Y., Becker, T., Beckmann, R. & Inada, T. 2022. Sensing of individual stalled 80S ribosomes by Fap1 for nonfunctional rRNA turnover. Mol Cell, 82, 3424–3437 e8.

Lu, P. D., Harding, H. P. & Ron, D. 2004. Translation reinitiation at alternative open reading frames regulates gene expression in an integrated stress response. J Cell Biol, 167, 27–33.

Lui, J., Castelli, L. M., Pizzinga, M., Simpson, C. E., Hoyle, N. P., Bailey, K. L., Campbell, S. G. & Ashe, M. P. 2014. Granules harboring translationally active mRNAs provide a platform for P-body formation following stress. Cell Rep, 9, 944–54.

Luo, Y., Na, Z. & Slavoff, S. A. 2018. P-Bodies: Composition, Properties, and Functions. Biochemistry, 57, 2424–2431.

Marton, M. J., Crouch, D. & Hinnebusch, A. G. 1993. GCN1, a translational activator of GCN4 in Saccharomyces cerevisiae, is required for phosphorylation of eukaryotic translation initiation factor 2 by protein kinase GCN2. Mol Cell Biol, 13, 3541–56.

Marton, M. J., Vazquez De Aldana, C. R., Qiu, H., Chakraburtty, K. & Hinnebusch, A. G. 1997. Evidence that GCN1 and GCN20, translational regulators of GCN4, function on elongating ribosomes in activation of eIF2alpha kinase GCN2. Mol Cell Biol, 17, 4474–89.

Masson, G. R. 2019. Towards a model of GCN2 activation. Biochem Soc Trans, 47, 1481–1488.

Morales-Polanco, F., Bates, C., Lui, J., Casson, J., Solari, C. A., Pizzinga, M., Forte, G., Griffin, C., Garner, K. E. L., Burt, H. E., Dixon, H. L., Hubbard, S., Portela, P. & Ashe, M. P. 2021. Core Fermentation (CoFe) granules focus coordinated glycolytic mRNA localization and translation to fuel glucose fermentation. iScience, 24, 102069.

Nanjaraj Urs, A. N., Kim, L. & Zaher, H. S. 2025. Insights into the role of collided ribosomes during the activation of the integrated stress response. Biochem Soc Trans, 53, 615–626.

Nanjaraj Urs, A. N., Lasehinde, V., Kim, L., Mcdonald, E., Yan, L. L. & Zaher, H. S. 2024. Inability to rescue stalled ribosomes results in overactivation of the integrated stress response. J Biol Chem, 300, 107290.

Natarajan, K., Meyer, M. R., Jackson, B. M., Slade, D., Roberts, C., Hinnebusch, A. G. & Marton, M. J. 2001. Transcriptional profiling shows that Gcn4p is a master regulator of gene expression during amino acid starvation in yeast. Mol Cell Biol, 21, 4347–68.

Nostramo, R., Varia, S. N., Zhang, B., Emerson, M. M. & Herman, P. K. 2016. The Catalytic Activity of the Ubp3 Deubiquitinating Protease Is Required for Efficient Stress Granule Assembly in Saccharomyces cerevisiae. Mol Cell Biol, 36, 173–83.

Nostramo, R., Xing, S., Zhang, B. & Herman, P. K. 2019. Insights into the Role of P-Bodies and Stress Granules in Protein Quality Control. Genetics, 213, 251–265.

Oltion, K., Carelli, J. D., Yang, T., See, S. K., Wang, H. Y., Kampmann, M. & Taunton, J. 2023. An E3 ligase network engages GCN1 to promote the degradation of translation factors on stalled ribosomes. Cell, 186, 346–362 e17.

Pizzinga, M., Bates, C., Lui, J., Forte, G., Morales-Polanco, F., Linney, E., Knotkova, B., Wilson, B., Solari, C. A., Berchowitz, L. E., Portela, P. & Ashe, M. P. 2019. Translation factor mRNA granules direct protein synthetic capacity to regions of polarized growth. J Cell Biol, 218, 1564–1581.

Pochopien, A. A., Beckert, B., Kasvandik, S., Berninghausen, O., Beckmann, R., Tenson, T. & Wilson, D. N. 2021. Structure of Gcn1 bound to stalled and colliding 80S ribosomes. Proc Natl Acad Sci U S A, 118.

Proud, C. G. 2005. eIF2 and the control of cell physiology. Semin Cell Dev Biol, 16, 3–12.

Sattlegger, E. & Hinnebusch, A. G. 2000. Separate domains in GCN1 for binding protein kinase GCN2 and ribosomes are required for GCN2 activation in amino acid-starved cells. EMBO J, 19, 6622–33.

Sattlegger, E. & Hinnebusch, A. G. 2005. Polyribosome binding by GCN1 is required for full activation of eukaryotic translation initiation factor 2alpha kinase GCN2 during amino acid starvation. J Biol Chem, 280, 16514–21.

Schneider, C. A., Rasband, W. S. & Eliceiri, K. W. 2012. NIH Image to ImageJ: 25 years of image analysis. Nat Methods, 9, 671–5.

Shenton, D., Smirnova, J. B., Selley, J. N., Carroll, K., Hubbard, S. J., Pavitt, G. D., Ashe, M. P. & Grant, C. M. 2006. Global translational responses to oxidative stress impact upon multiple levels of protein synthesis. J Biol Chem, 281, 29011–21.

Sheth, U. & Parker, R. 2003. Decapping and decay of messenger RNA occur in cytoplasmic processing bodies. Science, 300, 805–8.

Sikorski, R. S. & Hieter, P. 1989. A system of shuttle vectors and yeast host strains designed for efficient manipulation of DNA in Saccharomyces cerevisiae. Genetics, 122, 19–27.

Siu, F., Bain, P. J., Leblanc-Chaffin, R., Chen, H. & Kilberg, M. S. 2002. ATF4 is a mediator of the nutrient-sensing response pathway that activates the human asparagine synthetase gene. J Biol Chem, 277, 24120–7.

Standart, N. & Weil, D. 2018. P-Bodies: Cytosolic Droplets for Coordinated mRNA Storage. Trends Genet, 34, 612–626.

Sudhakar, A., Ramachandran, A., Ghosh, S., Hasnain, S. E., Kaufman, R. J. & Ramaiah, K. V. 2000. Phosphorylation of serine 51 in initiation factor 2 alpha (eIF2 alpha) promotes complex formation between eIF2 alpha(P) and eIF2B and causes inhibition in the guanine nucleotide exchange activity of eIF2B. Biochemistry, 39, 12929–38.

Swisher, K. D. & Parker, R. 2010. Localization to, and effects of Pbp1, Pbp4, Lsm12, Dhh1, and Pab1 on stress granules in Saccharomyces cerevisiae. PLoS One, 5, e10006.

Teixeira, D. & Parker, R. 2007. Analysis of P-body assembly in Saccharomyces cerevisiae. Mol Biol Cell, 18, 2274–87.

Teixeira, D., Sheth, U., Valencia-Sanchez, M. A., Brengues, M. & Parker, R. 2005. Processing bodies require RNA for assembly and contain nontranslating mRNAs. RNA, 11, 371–82.

Vattem, K. M. & Wek, R. C. 2004. Reinitiation involving upstream ORFs regulates ATF4 mRNA translation in mammalian cells. Proc Natl Acad Sci U S A, 101, 11269–74.

Vazquez De Aldana, C. R., Marton, M. J. & Hinnebusch, A. G. 1995. GCN20, a novel ATP binding cassette protein, and GCN1 reside in a complex that mediates activation of the eIF-2 alpha kinase GCN2 in amino acid-starved cells. EMBO J, 14, 3184–99.

Wang, C., Huang, Z., Du, Y., Cheng, Y., Chen, S. & Guo, F. 2010. ATF4 regulates lipid metabolism and thermogenesis. Cell Res, 20, 174–84.

Wang, Z., Zhang, C., Fan, C. & Liu, Y. 2023. Post-translational modifications in stress granule and their implications in neurodegenerative diseases. Biochim Biophys Acta Gene Regul Mech, 1866, 194989.

Wek, R. C. 2018. Role of eIF2alpha Kinases in Translational Control and Adaptation to Cellular Stress. Cold Spring Harb Perspect Biol, 10.

Wek, S. A., Zhu, S. & Wek, R. C. 1995. The histidyl-tRNA synthetase-related sequence in the eIF-2 alpha protein kinase GCN2 interacts with tRNA and is required for activation in response to starvation for different amino acids. Mol Cell Biol, 15, 4497–506.

Wheeler, J. R., Matheny, T., Jain, S., Abrisch, R. & Parker, R. 2016. Distinct stages in stress granule assembly and disassembly. Elife, 5.

Yang, R., Wek, S. A. & Wek, R. C. 2000. Glucose limitation induces GCN4 translation by activation of Gcn2 protein kinase. Mol Cell Biol, 20, 2706–17.

Youn, J. Y., Dyakov, B. J. A., Zhang, J., Knight, J. D. R., Vernon, R. M., Forman-Kay, J. D. & Gingras, A. C. 2019. Properties of Stress Granule and P-Body Proteomes. Mol Cell, 76, 286–294.

Zhu, M., Kuechler, E. R., Zhang, J., Matalon, O., Dubreuil, B., Hofmann, A., Loewen, C., Levy, E. D., Gsponer, J. & Mayor, T. 2020. Proteomic analysis reveals the direct recruitment of intrinsically disordered regions to stress granules in S. cerevisiae. J Cell Sci, 133.

