## Supplementary Table 1 for "The yeast Gcn2 protein kinase mediates P-body formation both through its activity with eIF2α and with non-canonical substrates"

| Yeast strains | | |
|---|---|---|
| Strain | Genotype | Source |
| W303-1A | *MATα ADE2 his3-11,15 leu2-3,112 trp1-1 ura3-1 can1-100* | Grant lab. |
| CY4322 | *DCP1-GFP::G418 PAB1-RFP::NAT* | Hoyle et al., (2007) |
| CY4479 | *DCP1-GFP::G418 PAB1-RFP::NAT gcn2::hphNT1* | This study. |
| CY4480 | *gcn2::hphNT1* | This study. |
| CY4573 | *CDC33-RFP::cloNAT DCP2-CFP::TRP ENO2-MS2:lox p[CP-GFP3 HIS3 CEN]* | Lui et al., (2014) |
| CY4574 | *lsm4ΔC::LEU2 edc3Δ::URA3 CDC33-RFP::cloNAT DCP2-CFP::TRP ENO2-MS2:lox p[CP-GFP3 HIS3 CEN]* | Lui et al., (2014) |
| CY4708 | *p[SUI2 LEU2] his3-11,15 leu2-3,112 trp1 ura3 sui2Δ* | Dever et al., (1992) |
| CY4709 | *p[SUI2-S51A LEU2] his3-11,15 leu2-3,112; trp1 ura3 sui2Δ* | Dever et al., (1992) |
| CY4743 | *p[SUI2 LEU2] his3-11,15 leu2-3,112 trp1 ura3 sui2Δ DCP1-GFP::kanMX* | This study. |
| CY4741 | *p[SUI2-S51A LEU2] his3-11,15 leu2-3,112 trp1 ura3 sui2Δ DCP1-GFP::kanMX* | This study. |
| CY4815 | *gcn1::hphNT1* | Grant lab. |
| CY4816 | *gcn3::hphNT1* | Grant lab. |
| CY4818 | *gcn20::hphNT1* | This study |
| CY4762 | *DCP1-GFP::G418 PAB1-RFP::NAT gcn1::hphNT1* | This study. |
| CY4727 | *DCP1-GFP::G418 PAB1-RFP::NAT gcn3::hphNT1* | This study |
| CY4763 | *DCP1-GFP::G418 PAB1-RFP::NAT gcn20::hphNT1* | This study. |
| CY4778 | *DCP1-GFP::G418 PAB1-RFP::NAT yar1::hphNT1* | This study. |
| CY4752 | *DCP1-GFP::G418 PAB1-RFP::NAT sap185::hphNT1* | This study |
| CY4756 | *DCP1-GFP::G418 PAB1-RFP::NAT smu2::hphNT1* | This study. |
| CY4750 | *DCP1-GFP::G418 PAB1-RFP::NAT ubp3::hphNT1* | This study. |
| CY4749 | *DCP1-GFP::G418 PAB1-RFP::NAT ssd1::hphNT1* | This study. |
| CY4757 | *DCP1-GFP::G418 PAB1-RFP::NAT fap1::hphNT1* | This study. |
| CY4759 | *DCP1-GFP::G418 PAB1-RFP::NAT mag2::hphNT1* | This study. |

| | | |
|---|---|---|
| CY4617 | *Dcp2-CFP::TRP1* | Hoyle et al., (2007) |
| CY4783 | *Dcp2-CFP::TRP1 Smu2-GFP::KanMX* | This study. |
| CY4784 | *Dcp2-CFP::TRP1 Fap1-GFP::KanMX* | This study. |
| CY4971 | *uS3-HA::hisMX6* | Li et al., (2022) |
| CY4793 | *uS3-HA::hisMX6 mag2::hphNT1* | This study. |
| CY4794 | *uS3-HA::hisMX6 smu2::hphNT1* | This study. |
| CY4953 | *smu2::hphNT1 p[DCP1-mCherry HIS3 CEN] p[SMU2*-GFP *TRP1 CEN]* | This study. |
| CY4954 | *smu2::hphNT1, p[DCP1-mCherry HIS3 CEN] p[SMU2-S32A-*GFP *TRP1 CEN]* | This study. |
| CY4179 | *DCP1-GFP::G418 DHH1-RFP::NAT* | Kershaw et al., (2021) |
