## Supplementary figures and images for "The yeast Gcn2 protein kinase mediates P-body formation both through its activity with eIF2α and with non-canonical substrates"

### Fig. S1

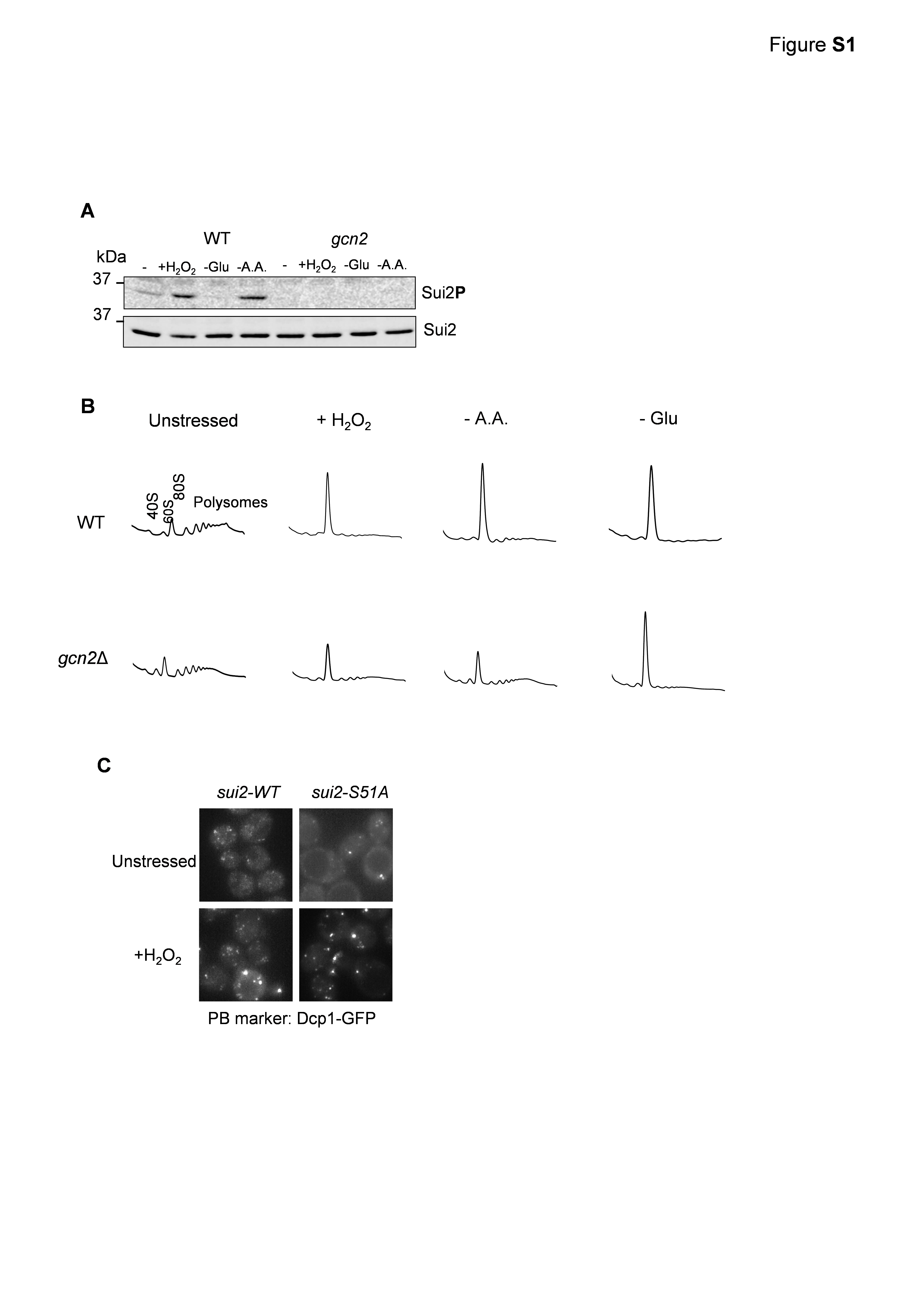
